# Comparable daughter radionuclide redistribution with superior tumor absorbed dose of the SSTR2 antagonist [^225^Ac]Ac-SSO110 versus [^225^Ac]Ac-DOTA-TATE

**DOI:** 10.64898/2026.03.16.711095

**Authors:** Prachi Desai, Marion Huber, Dennis Mewis, Nicolas Chouin, Manuel Sturzbecher-Hoehne, Germo Gericke, Anika Jaekel

## Abstract

It has been hypothesized that effective cellular internalization is required for the retention of ^225^Ac daughter radionuclides. The complex decay chain of ^225^Ac and recoil-mediated release of daughters, particularly ^213^Bi (half-life (t_1/2_) = 46 min), raise concerns about redistribution that may reduce tumor absorbed dose (TAD) and increase off-target radiation exposure. Because somatostatin receptor subtype 2 (SSTR2) antagonists such as SSO110 are not internalized, it has been proposed that the daughter radionuclides are less effectively retained compared to internalizing agonists such as DOTA-TATE. We therefore performed a direct and quantitative comparison of daughter radionuclide redistribution following administration of [^225^Ac]Ac-SSO110 and [^225^Ac]Ac-DOTA-TATE.

**Methods:** Biodistribution and ^213^Bi redistribution were evaluated in Balb/c nude mice bearing NCI-H69 small cell lung cancer xenografts. Repeated gamma counting combined with bi-exponential modeling was used to quantify ^225^Ac and ^213^Bi activity in tumor, blood, bone marrow, kidneys, liver, and intestines up to 96 h post-injection. TAD was calculated with and without accounting for experimentally-derived ^213^Bi redistribution. Real-time in vitro binding assays were conducted to characterize cellular retention of [^225^Ac]Ac-SSO110.

**Results:** [^225^Ac]Ac-SSO110 demonstrated higher tumor uptake and prolonged retention compared with [^225^Ac]Ac-DOTA-TATE, resulting in a 1.9-fold higher tumor-to-kidney ratio at 96 h and a 2.8-fold higher TAD. Redistribution of ^213^Bi from tumor was minimal and comparable between agonist and antagonist, with maximum tumor loss of 3.5% for [^225^Ac]Ac-SSO110 and 2% for [^225^Ac]Ac-DOTA-TATE. Accounting for daughter redistribution reduced TAD by less than 5% for both radioconjugates. No sustained ^213^Bi accumulation was observed in blood, kidneys, or liver, and only minimal activity was detected in bone marrow and intestines. Real-time binding studies demonstrated sustained cell-associated β^-^ signal following incubation with [^225^Ac]Ac-SSO110.

**Conclusion:** Receptor-mediated internalization is not required for effective retention of ^225^Ac daughter radionuclides. Despite negligible internalization, [^225^Ac]Ac-SSO110 achieved superior TAD and higher tumor-to-kidney ratio without increased daughter redistribution compared with the internalizing agonist [^225^Ac]Ac-DOTA-TATE. These findings question the necessity of internalization for daughter retention and support further evaluation of antagonist-based ^225^Ac radioligand therapy.

## INTRODUCTION

Targeted radionuclide therapy directed against SSTR2 with the β-emitting SSTR2 agonist [^177^Lu]Lu-DOTA-TATE has become an established treatment modality for neuroendocrine tumors (*1,2*). However, durable tumor control remains incomplete, particularly in patients with high tumor burden and heterogenous receptor expression (*3*). Increasing tumor absorbed dose while maintaining acceptable organ exposure therefore remains a central objective in the ongoing development of SSTR2-directed radioligand therapy.

SSTR2 antagonists, such as SSO110, have emerged as next-generation ligands, as they recognize a broader spectrum of receptor conformations and bind to a greater number of binding sites. SSO110 achieved higher tumor accumulation than DOTA-TATE (*4*–*7*) in preclinical models. Consistent with these findings, clinical studies demonstrated that [^177^Lu]Lu-SSO110 delivers higher TAD and achieves higher tumor-to-organ ratios than [^177^Lu]Lu-DOTA-TATE (*8*), supporting the therapeutic potential of SSTR2 antagonist-based targeting.

Combining the tumor-targeting advantage of SSTR2 antagonists with high linear energy transfer (LET) α-emitters represents a promising strategy to enhance therapeutic efficacy. ^225^Ac is particularly attractive for SSTR2-directed targeted alpha therapy because its physical t_1/2_ of 9.9 days aligns well with the prolonged tumor residence time of SSO110, enabling sustained irradiation of tumor tissue. The decay of ^225^Ac results in the emission of four α-particles with energies ranging from 5.8 MeV to 8.4 MeV, which are highly effective in inducing double-strand DNA breaks. The combination of long t_1/2_ with sequential α-decays enables sustained and concentrated energy deposition over short path lengths, thereby creating irreparable DNA damage while minimizing crossfire irradiation to surrounding healthy tissues (*9*). In contrast, the relatively low LET and millimeter-range tissue penetration of β-particles may limit their effectiveness to induce lethal DNA damage at the single-cell level (*10*), particularly in settings of heterogeneous target expression.

However, the same decay cascade that underlies the therapeutic potential of ^225^Ac also introduces a key challenge: namely recoil-induced de-chelation and potential redistribution of daughter radionuclides. The recoil energy generated during α-decay (∼ 100 to 200 keV) is sufficient to disrupt chemical bonds (*11*), thereby releasing daughter radionuclides from the chelator. Because each daughter radionuclide possesses distinct chemical properties and biodistribution characteristics, their release may decrease TAD and increase off-target radiation exposure.

The short-lived daughters ^221^Fr (t_1/2_=4.9 min) and ^217^At (t_1/2_=32 ms) are generally assumed to decay near the site of the parent radionuclide due to the limited time available for migration (*12*). In contrast, the longer-lived daughter ^213^Bi (t_1/2_=46 min) may have sufficient time to redistribute before decay, thereby possibly diminishing tumor absorbed dose and increasing radiation burden to healthy tissues (*13*).

It has been widely proposed that rapid radioligand-receptor internalization limits systemic redistribution of ^225^Ac daughter radionuclides by trapping them intracellularly (*14*). According to this paradigm, internalizing agonists such as DOTA-TATE would be expected to retain ^225^Ac daughter radionuclides more efficiently than non-internalizing antagonists such as SSO110. In preclinical models, [^225^Ac]Ac-SSO110 demonstrated greater tumor growth inhibition than the internalizing agonist [^225^Ac]Ac-DOTA-TATE, without evidence of increased organ toxicity (*15*). These findings challenge the prevailing internalization-retention paradigm and highlight the limited experimental evidence directly comparing daughter radionuclide redistribution between antagonist- and agonist-based ^225^Ac-labeled radiopharmaceuticals.

In the present study, we performed a direct and quantitative evaluation of daughter radionuclide redistribution following administration of [^225^Ac]Ac-SSO110 and [^225^Ac]Ac-DOTA-TATE to address these mechanistic and safety questions. Experimentally derived ^213^Bi time-activity curves were generated in tumor tissue and selected organs to characterize biodistribution. Tumor uptake data were subsequently integrated to determine the impact of daughter migration on TAD. In addition, real-time cellular binding analyses were conducted to characterize the cellular retention behavior of [^225^Ac]Ac-SSO110. Through this integrated approach, we aimed to determine whether antagonist-based targeting increases daughter release compared with internalizing agonists. A clear understanding of daughter redistribution is essential for the rational clinical development of ^225^Ac-labeled SSTR2-targeted radioligand therapy.

## MATERIALS AND METHODS

### Radiochemistry

#### [^177^Lu]Lu-SSO110

In a microtube [^177^Lu]LuCl_3_ (ITM/Eckert & Ziegler, 0.04 M HCl) was mixed with radiolabeling buffer (pH 5.1) and SSO110 (dissolved in 0.1% TFA). The reaction mixture was incubated at 90 °C for 30 minutes. At the end of the incubation, 4 mM DTPA in 0.9% NaCl was added at room temperature to chelate unbound ^177^Lu. Final solutions were diluted in 0.9% NaCl supplemented with 5 mg/mL ascorbic acid. The final specific activity ranged from 20 MBq/µg to 59 MBq/µg.

#### [^225^Ac]Ac-SSO110 and [^225^Ac]Ac-DOTA-TATE

In an Eppendorf tube, [^225^Ac]AcCl_3_ (ITM, 0.04 M HCl) was mixed with radiolabeling buffer (pH 8.1) and SSO110 or DOTA-TATE and heated at 80 °C for 20-30 minutes. The reaction mixture was cooled to room temperature, followed by addition of a stabilizing solution containing 100 mg/mL sodium ascorbate and 0.1 mg/mL DTPA. The final specific activity was 90 kBq/µg for both radioconjugates.

Radiolabeling incorporation was assessed by iTLC for all radioconjugates, and radiochemical purity was confirmed by HPLC on representative batches. All preparations demonstrated ≥ 95% incorporation and, where evaluated by HPLC, radiochemical purity was ≥ 95%.

### Biodistribution and quantitative assessment of ^213^Bi redistribution

The biodistribution of [^225^Ac]Ac-SSO110 and [^225^Ac]Ac-DOTA-TATE was evaluated in Balb/c nude mice bearing NCI-H69 SCLC tumor xenografts. Mice received a single intravenous injection of 90 kBq [^225^Ac]Ac-SSO110 and [^225^Ac]Ac-DOTA-TATE.

At predefined time points post-injection, blood, tumor, bone marrow, intestines, kidneys, and liver were collected and weighed. Radioactivity was counted using a gamma counter with a 400-480 keV energy window specific for ^213^Bi. Individual ^213^Bi counts were recorded every 13 minutes for 20 hours and fitted using a bi-exponential function. Counts obtained before secular equilibrium were used to quantify loss or excess of ^213^Bi relative to ^225^Ac. After secular equilibrium, ^213^Bi counts reflect ^225^Ac activity and were decay-corrected to derive ex vivo biodistribution (%IA/g; mean ± SD). Tumor-to-kidney ratios were calculated to evaluate tumor-to-organ distribution over time.

All animal experiments were approved by the relevant institutional and national authorities for animal welfare and were conducted in accordance with applicable guidelines.

### Tumor absorbed dose calculations

Tumor time–activity curves were fitted using mono- or bi-exponential models using GraphPad PRISM 7 (GraphPad Software Inc., San Diego, CA, USA). The TAD was calculated according to the MIRD formalism (*16*) under two scenarios. In the first scenario, no redistribution of daughter radionuclides after ^225^Ac decay was assumed, and TAD was calculated by multiplying the time-integrated activity per gram of tumor (expressed in Bq*s/g) by the total energy emitted via α-particles (27.52 MeV). In the second scenario, ^213^Bi loss from the tumor was considered based on the experimentally derived ^213^Bi time-activity curves up to 96 h. TAD was calculated as the sum of the time-integrated activity per gram of tumor (expressed in Bq*s/g), derived for ^225^Ac, multiplied by the energy emitted via α-particles for ^225^Ac, ^221^Fr and ^217^At (19.20 MeV), and the time-integrated activity per gram of tumor (expressed in Bq*s/g), derived for ^213^Bi, multiplied by the energy emitted via α-particles for ^213^Bi and ^213^Po (8.32 MeV).

### Cellular binding and affinity

The cellular binding characteristics and affinity of [^225^Ac]Ac-SSO110 and [^177^Lu]Lu-SSO110 were determined using the saturation binding assay and real-time binding analyses.

For the saturation binding assay, SSTR2-positive AR42J cells were incubated for 4 h at 37 °C with increasing concentrations of [^177^Lu]Lu-SSO110 (0 to 1.5 nM) and [^225^Ac]Ac-SSO110 (0 to 3 nM). Non-specific binding was determined in the presence of a 100-fold excess of non-radiolabeled SSO110. At the end of the incubation period, supernatants containing unbound radioligand were collected, and cell-associated radioactivity was recovered by lysis with 1 M NaOH for 5 minutes. Bound radioactivity was quantified using a gamma counter. Because total binding exceeded 10% of the added radioligand, ligand depletion was accounted for during nonlinear regression using the “One site - Total binding’’ model in GraphPad Prism (v10.3.1. Equilibrium dissociation constants (K_D_) were derived from the fitted curves.

Real-time binding of [^225^Ac]Ac-SSO110 and [^177^Lu]Lu-SSO110 was performed using LigandTracer^®^ White (Ridgeview Instruments AB, Sweden). AR42J (SSTR2-positive) and PC-3 (SSTR2-negative control; only tested with [^225^Ac]Ac-SSO110) cells were seeded at 2×10^6^ in petri dishes. Radioconjugates were tested at concentrations 0.18 to 0.53 nM for [^177^Lu]Lu-SSO110 and 1.7 to 3.4 nM for [^225^Ac]Ac-SSO110 to obtain association curves. After reaching the plateau, the radioactive medium was replaced with fresh complete medium to determine the dissociation. All measurements were performed at room temperature. Specificity of the [^225^Ac]Ac-SSO110 signal in AR42J cells was confirmed by blocking with excess non-radiolabeled SSO110. Binding data were evaluated using TraceDrawer (1.9.2, Ridgeview Instruments AB, Sweden) using a 1:1 interaction model.

## RESULTS

### Higher tumor uptake and prolonged tumor retention of [^225^Ac]Ac-SSO110 compared with [^225^Ac]Ac-DOTA-TATE

Biodistribution was evaluated in Balb/c nude mice bearing NCI-H69 xenografts following administration of [^225^Ac]Ac-SSO110 and [^225^Ac]Ac-DOTA-TATE. This model was selected because its moderate and heterogenous SSTR2 expression reflects clinically-relevant receptor levels (*17,18*). Both compounds rapidly cleared from blood, with radioactivity decreasing to below the limit of detection by 48 h post injection (Fig. 1). Tumor uptake of [^225^Ac]Ac-SSO110 exceeded that of [^225^Ac]Ac-DOTA-TATE at all evaluated time points. Peak tumor accumulation of [^225^Ac]Ac-SSO110 was observed at 4 h post injection (6.8% IA/g), whereas [^225^Ac]Ac-DOTA-TATE reached its maximum tumor uptake earlier at 0.5 h (5.0% IA/g) and declined thereafter. At later time points, tumor activity concentrations remained higher for [^225^Ac]Ac-SSO110, indicating prolonged tumor retention. High renal uptake was observed early after injection, consistent with renal excretion and partial reabsorption, reaching 23.7% IA/g for [^225^Ac]Ac-SSO110 and 19% IA/g for [^225^Ac]Ac-DOTA-TATE at 0.5 h post injection. Uptake in liver, intestines, and bone marrow remained below 2.5% IA/g throughout the observation period. Radioactivity in all organs declined over time.

**Figure 1.**
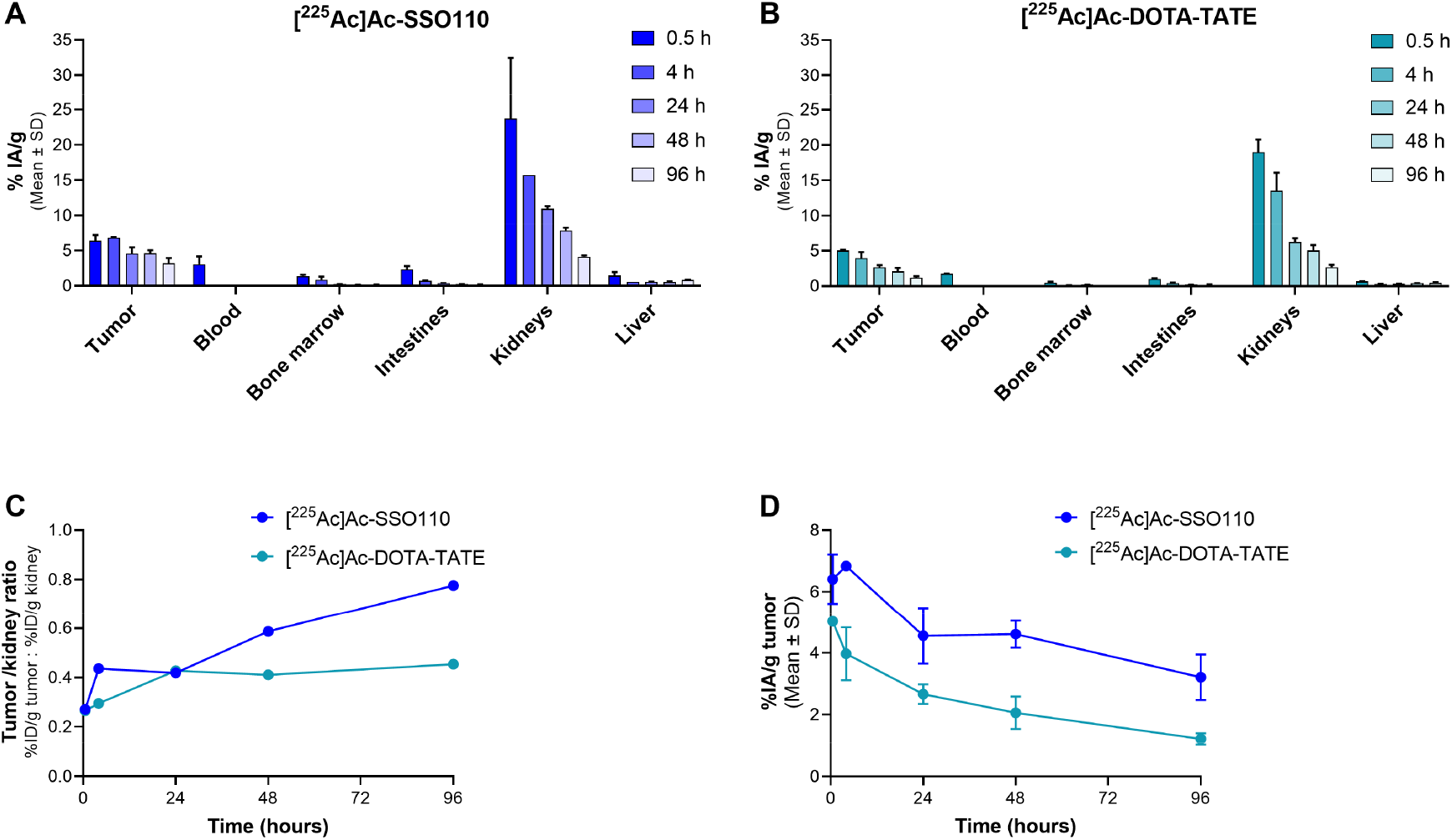
Biodistribution of ^225^Ac-labeled SSO110 and DOTA-TATE up to 96 h post-injection. % IA/g of selected organs following administration of (A) 90 kBq [^225^Ac]Ac-SSO110 (n=2) and (B) [^225^Ac]Ac-DOTA-TATE (n=2). (C) Tumor-to-kidney ratios and (D) tumor area under the curve of [^225^Ac]Ac-SSO110 and [^225^Ac]Ac-DOTA-TATE. Data are presented as mean ± SD.

Tumor-to-kidney ratios increased over time for both compounds and were consistently higher for [^225^Ac]Ac-SSO110, reaching a maximum 1.9-fold higher ratio than [^225^Ac]Ac-DOTA-TATE at 96 h post injection (Fig. 1C). Integration of the tumor time-activity curves yielded a 2.0-fold greater tumor area under the curve (AUC) for [^225^Ac]Ac-SSO110 (Fig. 1D).

Collectively, these findings demonstrate enhanced tumor exposure and favorable tumor-to-kidney ratio for [^225^Ac]Ac-SSO110 compared with [^225^Ac]Ac-DOTA-TATE.

### Minor loss of ^213^Bi from tumor after administration of [^225^Ac]Ac-SSO110 and [^225^Ac]Ac-DOTA-TATE with minimal impact on tumor absorbed dose

Redistribution of ^213^Bi, the third daughter radionuclide of ^225^Ac, was evaluated in tumor tissue by comparing measured ^213^Bi activity with the corresponding ^225^Ac activity derived from the biodistribution study. Deviations of ^213^Bi activity relative to ^225^Ac were interpreted as evidence of ^213^Bi loss or gain.

In tumors, a small early loss of ^213^Bi relative to ^225^Ac was observed for both radioconjugates. The maximum relative loss occurred at 0.5 h post injection, reaching 3.5% for [^225^Ac]Ac-SSO110 and 2% for [^225^Ac]Ac-DOTA-TATE. This loss decreased over time, with ^213^Bi loss falling below 1% from 24 h to 96 h p.i (Fig. 2A).

**Figure 2.**
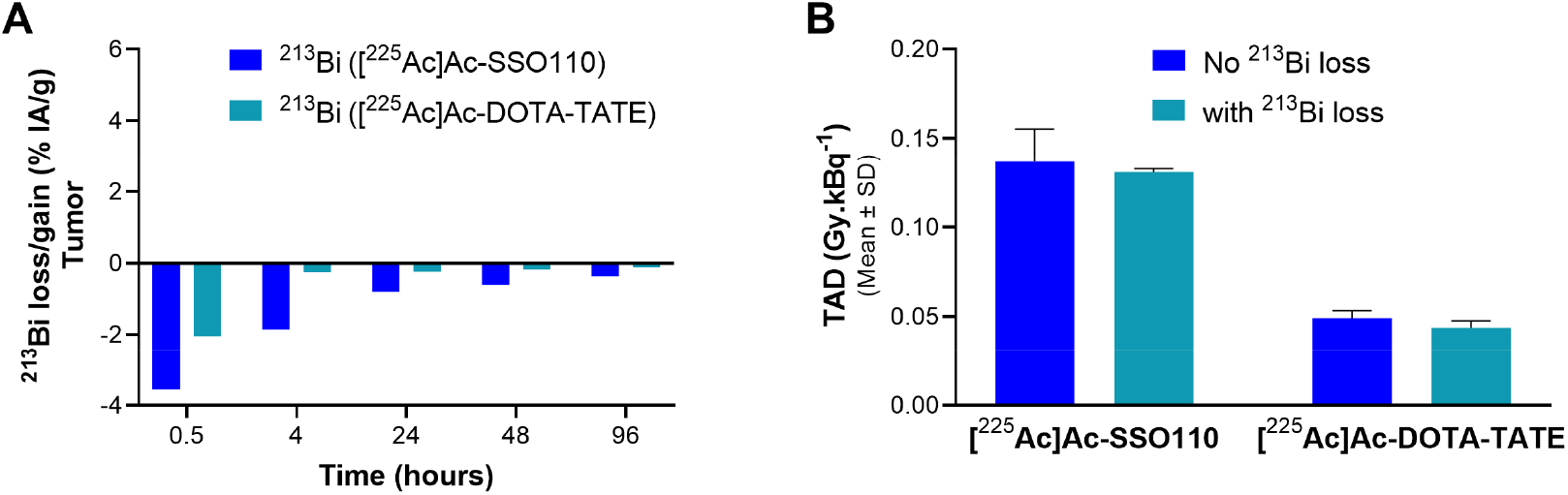
^213^Bi redistribution of ^225^Ac-labeled SSO110 and DOTA-TATE in tumor tissue and its impact on TAD. (A) Relative ^213^Bi loss or gain compared with ^225^Ac, represented as %IA/g for tumor. Positive values indicate ^213^Bi gain, and negative values indicate ^213^Bi loss relative to ^225^Ac. (B) TAD calculated with and without correction for measured ^213^Bi loss. Data are presented as mean ± SD.

To determine the dosimetric relevance of this redistribution, TAD was calculated with and without correction for measured ^213^Bi loss. ^213^Bi redistribution from tumor resulted in only minor reductions in TAD, with a maximum decrease of 4.5% for [^225^Ac]Ac-SSO110 and 3.9% for [^225^Ac]Ac-DOTA-TATE (Fig. 2B). Importantly, [^225^Ac]Ac-SSO110 delivered a 2.8-fold higher TAD compared with [^225^Ac]Ac-DOTA-TATE.

### Similar ^213^Bi redistribution in healthy organs for [^225^Ac]Ac-SSO110 and [^225^Ac]Ac-DOTA-TATE

Redistribution of ^213^Bi in non-target organs was evaluated in blood, bone marrow, kidneys, liver, and intestines by comparing measured ^213^Bi activity with corresponding ^225^Ac activity.

In blood, an early loss of ^213^Bi relative to ^225^Ac was observed at 0.5 h post injection for both radioconjugates. At later time points, ^213^Bi activity in blood was below the limit of detection (Fig. 3A). In bone marrow, variable relative gain was observed across time points for both compounds (Fig. 3B). However, absolute activity concentrations remained consistently below 0.5% IA/g, as shown in the biodistribution data (Fig. 1).

**Figure 3.**
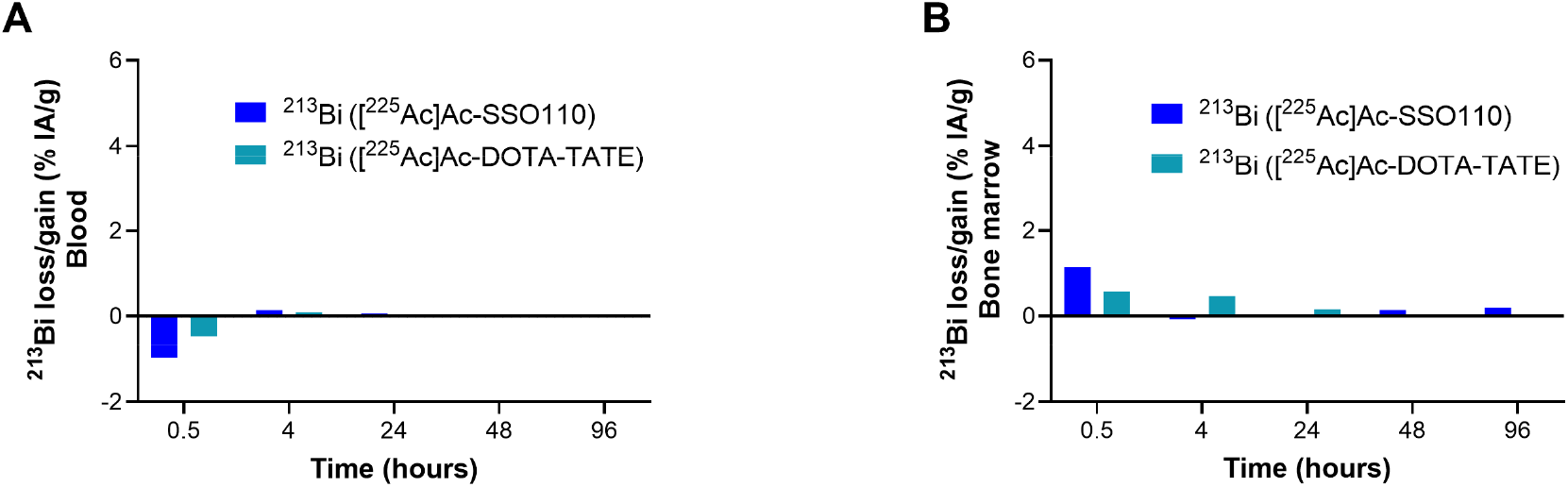
^213^Bi redistribution of ^225^Ac-labeled SSO110 and DOTA-TATE in blood and bone marrow. ^213^Bi loss or gain relative to ^225^Ac represented as % IA/g for (A) blood, and (B) bone marrow. Positive Y-axis denotes ^213^Bi gain, and negative Y-axis denotes ^213^Bi loss relative to ^225^Ac.

In kidneys, liver, and intestines, a relative gain of ^213^Bi compared with ^225^Ac was detected at 0.5 h post injection for both radioconjugates (Fig. 4). At subsequent time points, ^213^Bi activity remained below corresponding ^225^Ac activities. In the intestines, some variability in relative ^213^Bi gain persisted beyond 0.5 h for both compounds, while absolute ^213^Bi activities remained below 0.5% IA/g (Fig. 4C).

**Figure 4.**
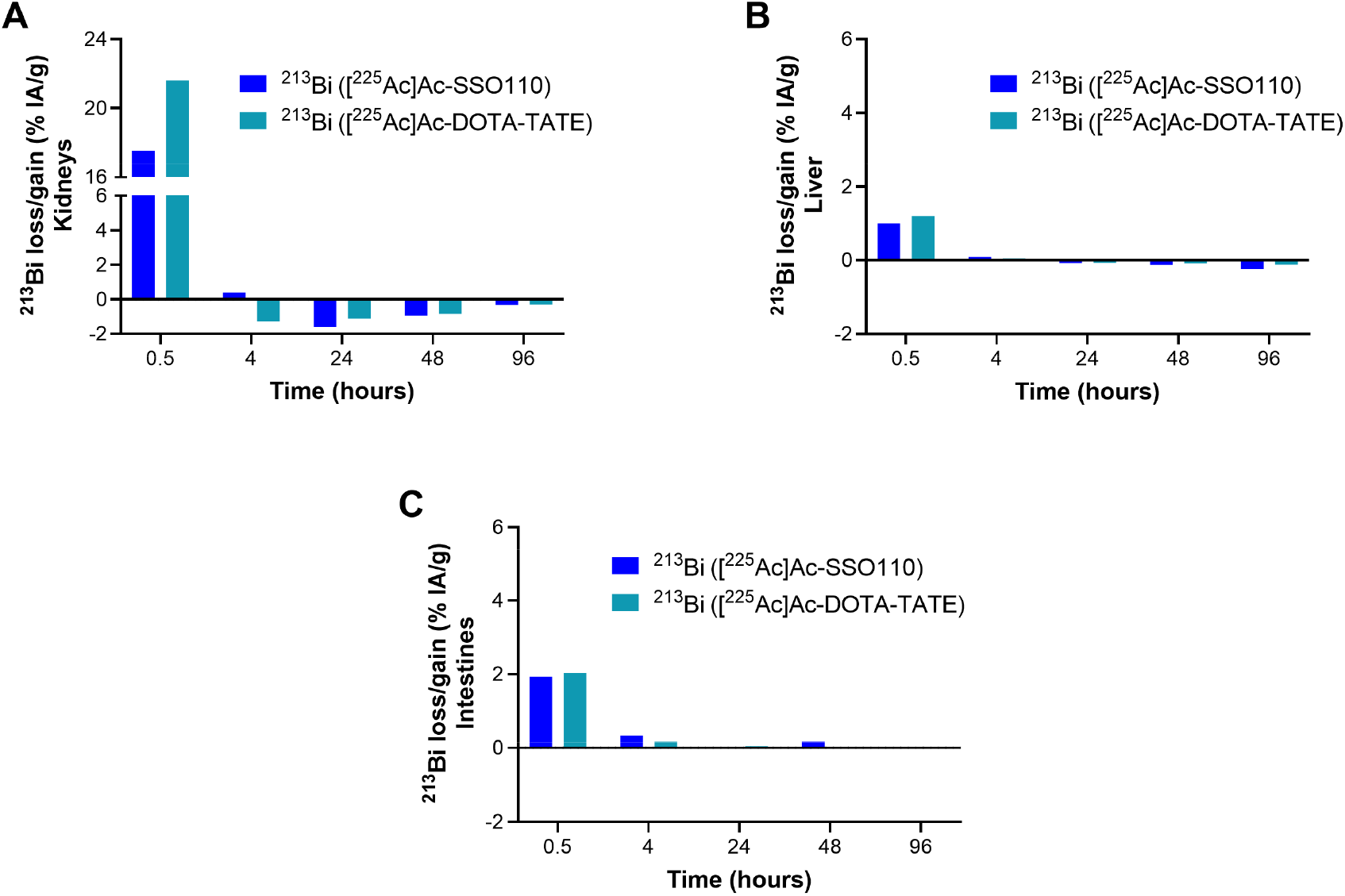
^213^Bi redistribution of ^225^Ac-labeled SSO110 and DOTA-TATE in kidneys, liver, and intestines. ^213^Bi loss or gain relative to ^225^Ac represented as % IA/g for (A) kidneys, (B) liver, and (C) intestines. Positive Y-axis denotes ^213^Bi gain, and negative Y-axis denotes ^213^Bi loss relative to ^225^Ac.

Overall, patterns of ^213^Bi redistribution in healthy organs were comparable between [^225^Ac]Ac-SSO110 and [^225^Ac]Ac-DOTA-TATE. No accumulation of ^213^Bi was observed in blood, kidneys, or liver, and only minimal activity was detected in intestines and bone marrow.

### Real-time binding studies indicate sustained cell-associated β^-^ signal following incubation with [^225^Ac]Ac-SSO110

To further investigate cellular retention, real-time binding kinetics were analyzed in SSTR2-positive AR42J cells and SSTR2-negative PC-3 cells, including specificity controls. Binding of [^225^Ac]Ac-SSO110 was monitored using a β-detection LigandTracer^®^ White.

In AR42J cells, the β^-^ signal increased gradually during the association phases and reached a plateau over several hours. During the dissociation phase, no measurable decrease in signal was observed within the experimental timeframe, indicating sustained cell-associated β^-^ signal (Fig. 5A). Specificity of the signal was confirmed by the absence of measurable binding in SSTR2-negative PC-3 cells and by competitive blocking with excess non-radioactive SSO110 in AR42J cells (Fig. 5B, C). For comparison, real-time binding of [^177^Lu]Lu-SSO110 in AR42J cells demonstrated slow but measurable dissociation during washout (Fig. S2).

**Figure 5.**
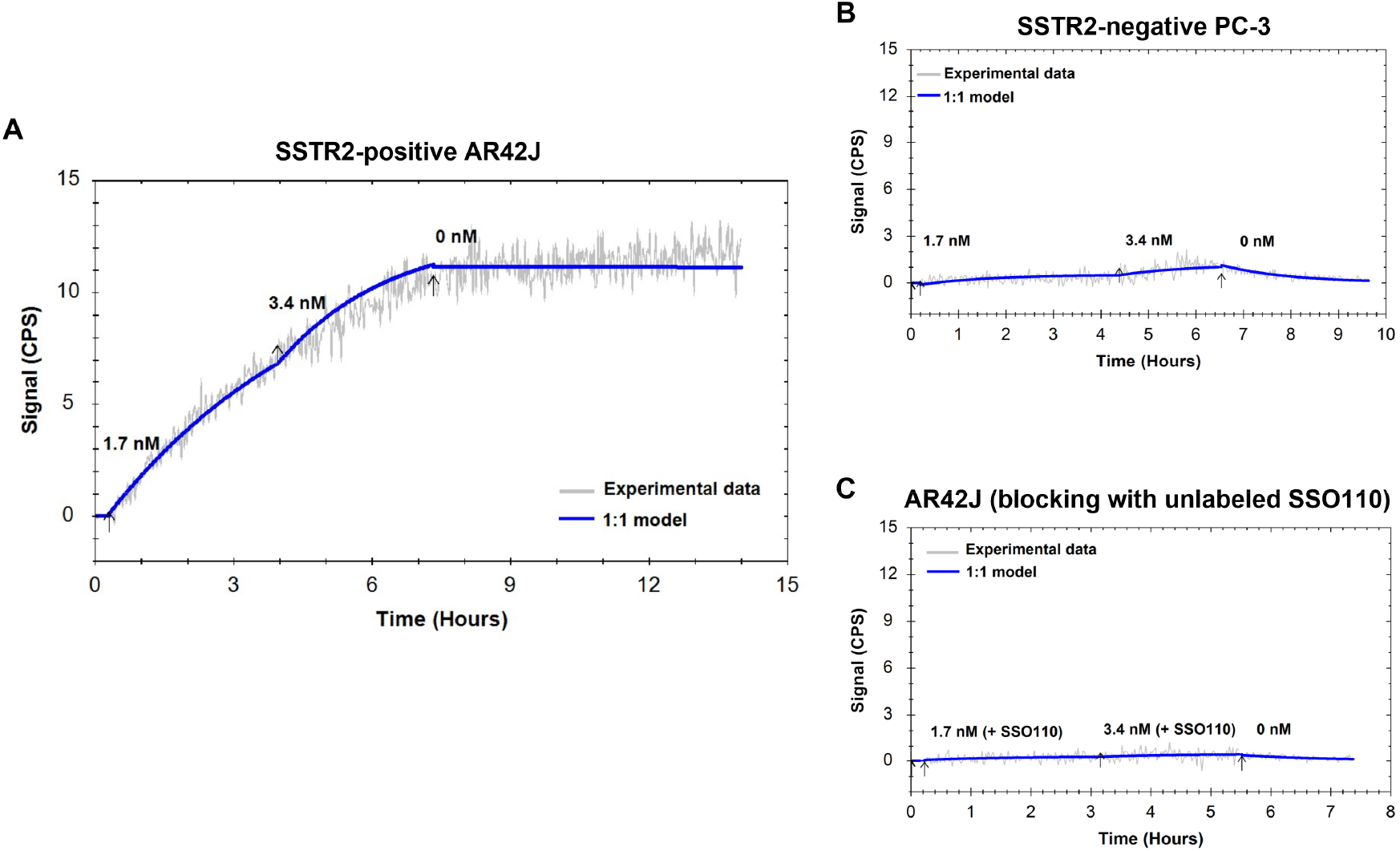
Real-time cellular binding of [^225^Ac]Ac-SSO110. Association curves following incubation with 1.7 nM and 3.4 nM [^225^Ac]Ac-SSO110 and subsequent dissociation in (A) high SSTR2-expressing AR42J cells, (B) low-to-negative SSTR2-expressing PC-3 cells, and (C) high SSTR2-expressing AR42J cells in the presence of excess non-radiolabeled SSO110. Arrows indicate the time points of compound or medium addition.

Importantly, the cellular affinity determined using saturation binding assays were comparable for [^225^Ac]Ac-SSO110 and [^177^Lu]Lu-SSO110 (Table 1), indicating that substitution of the radiometal did not alter receptor affinity.

**Table 1.** Cellular affinity of [^225^Ac]Ac-SSO110 and [^177^Lu]Lu-SSO110 determined by saturation binding assays.

| Radiopeptides | $k_D$ (nM) |
| --- | --- |
| [$^{225}\text{Ac}$]Ac-SSO110 | 0.42 |
| [$^{177}\text{Lu}$]Lu-SSO110 | 0.29 |

Overall, these data support sustained cellular retention of β^-^-derived daughter activity following incubation with [^225^Ac]Ac-SSO110.

## DISCUSSION

This study provides a direct head-to-head comparison of daughter radionuclide redistribution between an internalizing SSTR2 agonist, [^225^Ac]Ac-DOTA-TATE, and a non-internalizing SSTR2 antagonist, [^225^Ac]Ac-SSO110. Contrary to the prevailing assumption that receptor-mediated internalization is required to retain ^225^Ac recoil daughters within the tumor, both radioconjugates demonstrated comparable ^213^Bi redistribution profiles. Tumor-associated ^213^Bi loss was minimal for both compounds with negligible impact of daughter redistribution on tumor dosimetry. Furthermore, no sustained retention of ^213^Bi was observed in blood, kidneys, or liver, and only minimal accumulation was detected in intestines and bone marrow.

Despite absence of receptor-mediated internalization, [^225^Ac]Ac-SSO110 achieved higher tumor uptake, prolonged retention, an up-to-1.9-fold higher tumor-to-kidney ratio, and a 2.8-fold higher TAD compared with [^225^Ac]Ac-DOTA-TATE. The maximum early ^213^Bi loss observed in tumor after [^225^Ac]Ac-SSO110 and [^225^Ac]Ac-DOTA-TATE injections decreased over time and remained below 1 % beyond 24 h, and the total reduction in TAD remained below 5%. These data indicate that lack of internalization does not translate into increased macroscopic daughter radionuclide escape.

Real-time cellular binding studies further supported restricted daughter redistribution of [^225^Ac]Ac-SSO110. Sustained cell-associated β^-^ signal was observed after incubation of SSTR2-positive cells with [^225^Ac]Ac-SSO110, whereas [^177^Lu]Lu-SSO110 showed classical dissociation kinetics. As both radioconjugates demonstrated comparable receptor affinities, the sustained β^-^ signal is unlikely to be explained by differences in receptor binding. Although the underlying mechanisms were not directly investigated, the short recoil range of daughter radionuclides (*19*) together with limited diffusion at the plasma membrane and within the tumor microenvironment may contribute to restricted redistribution. These considerations are consistent with the sustained cell-associated β^-^ signal observed in vitro.

In healthy organs, no sustained ^213^Bi accumulation was observed in blood, kidney, or liver. Both radioconjugates cleared rapidly from circulation, with blood activity falling below detection by 48 h, thereby limiting systemic exposure time for daughter redistribution. Early deviations at 0.5 h, characterized by relative ^213^Bi loss in blood and transient gain in kidneys, liver, and intestines, likely reflect redistribution of circulating daughters or daughter-DTPA-complexes present at the time of injection. Importantly, no relevant ^213^Bi gain relative to ^225^Ac was detected in healthy organs at later time points. Absolute activity concentrations in intestines and bone marrow remained low throughout the observation period. Although relative fluctuations were observed in bone marrow, activities remained below 0.5% IA/g and did not show progressive accumulation.

The interpretation of these early non-equilibrium effects is supported by daughter-specific biodistribution studies. Pure ^221^Fr preferentially accumulates in kidney, salivary glands, and small intestine, whereas ^213^Bi exhibits renal and hepatic tropism (*20,21*). Thus, transient early uptake in clearance organs most likely reflects intrinsic daughter-organ affinity rather than recoil-mediated tumor release.

The magnitude of daughter contribution to organ dose appears to vary across types of targeting ligands. In a PSMA-targeted ^225^Ac small-molecule radioligand study, Wurzer et al. (*22*) observed no detectable redistribution of ^213^Bi from tumor. However, incorporation of daughter contributions resulted in moderate increases in kidney (1.1 to 1.3-fold) and salivary gland (1.5 to 2.5-fold) absorbed-dose estimates, due to early non-equilibrium uptake. In contrast, substantially larger effects have been reported for long-circulating antibody-based ^225^Ac constructs, where redistribution of daughters generated in circulation persisted more than 10 days and was associated with markedly increased kidney and salivary gland doses (Schatz et al., EANM 2024, oral presentation, unpublished data). In the present study, only minimal tumor ^213^Bi loss and transient early redistribution to kidneys were observed, without evidence of progressive accumulation in healthy organs. Although healthy organ absorbed doses were not formally calculated, the redistribution pattern observed here more closely resembles that reported for rapidly clearing small-molecule radioligands (*22*) than for long-circulating antibody-based constructs. Thus, the relatively rapid blood clearance of peptide-based radiopharmaceuticals may therefore represent a safety advantage compared with compounds characterized by prolonged systemic exposure.

Together, these observations suggest that daughter redistribution at the organ level is primarily influenced by ligand pharmacokinetics, receptor affinity, and the site of radionuclide decay, rather than internalization.

Redistribution of ^221^Fr, the first daughter in the ^225^Ac decay chain, was assessed exploratorily and definitive conclusions could not be drawn due to the short half-life of ^221^Fr and its fast equilibrium kinetics. Given the established salivary tropism of free ^221^Fr, further dedicated investigation is warranted. Salivary glands were not collected in this study and therefore potential accumulation of ^213^Bi or ^221^Fr in this organ cannot be excluded.

Our findings are consistent with clinical imaging data from the ACTION-1 study, which demonstrated limited daughter redistribution following administration of [^225^Ac]Ac-DOTA-TATE (RAYZ101) (*23*).

## CONCLUSION

This head-to-head comparison demonstrates that receptor-mediated internalization is not required for effective daughter retention within rapidly clearing peptide-based ^225^Ac therapies. Despite absence of internalization, the high-affinity antagonist SSO110 achieved higher tumor uptake, prolonged retention, and substantially greater tumor-absorbed dose with minimal daughter redistribution. These data support the concept that ligand pharmacokinetics and receptor affinity outweigh internalization kinetics as determinants of macroscopic daughter retention and provide a mechanistic basis for the superior therapeutic profile observed for [^225^Ac]Ac-SSO110.

## ACKNOWLEDGMENTS

The authors thank Chelatec for conducting the in vivo studies. The authors are grateful to Shelby Hartzell for language editing and review.

