## Supplemental Data for "Comparable daughter radionuclide redistribution with superior tumor absorbed dose of the SSTR2 antagonist [^225^Ac]Ac-SSO110 versus [^225^Ac]Ac-DOTA-TATE"

**Supplemental methods**

**Quantification of $^{225}$Ac and $^{213}$Bi activities at time of death**

To determine $^{225}$Ac and $^{213}$Bi activities in tissues at the time of death, repeated gamma counting was performed using the $^{213}$Bi energy window (400-480 keV). Samples were measured for 30s at 13-minute intervals. Activities of $^{225}$Ac and $^{213}$Bi were separated using a bi-exponential decay model as previously described (*1*).

Values of repeated measurements were represented as function of time (Fig. S1) and were fitted according to the following equation.

$$\frac{CPM_{EW\ 213Bi}\ (t)}{CF213_{Bi}} = A_{total,213_{Bi}}(t) = A^0 225_{Ac} \times e^{-\lambda 225_{Ac} t} + A^0 213_{Bi} \times e^{-\lambda 213_{Bi} t}$$

Where, $CPM_{EW\ 213Bi}\ (t)$ represents the gamma counter measurement at time t using the $^{213}$Bi energy window (EW), $CF213_{Bi}$ represents the calibration factor for $^{213}$Bi, and $\lambda 225_{Ac}$ and $\lambda 213_{Bi}$ represents the physical decay constants for $^{225}$Ac and $^{213}$Bi, respectively.

**Supplemental figures**

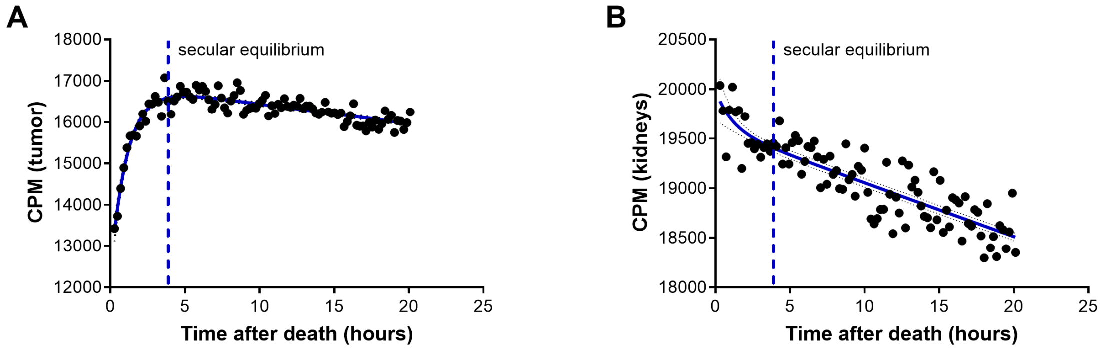


**Figure S1.** Exemplary time-dependent gamma counting following tissue collection. Counts per minute (CPM) measured in the $^{213}$Bi energy window as a function of time after death in (A) tumor, and (B) kidneys, 4 h after injection of [$^{225}$Ac]Ac-SSO110. The dashed line indicates the time point at which secular equilibrium between $^{225}$Ac and $^{213}$Bi is reached.

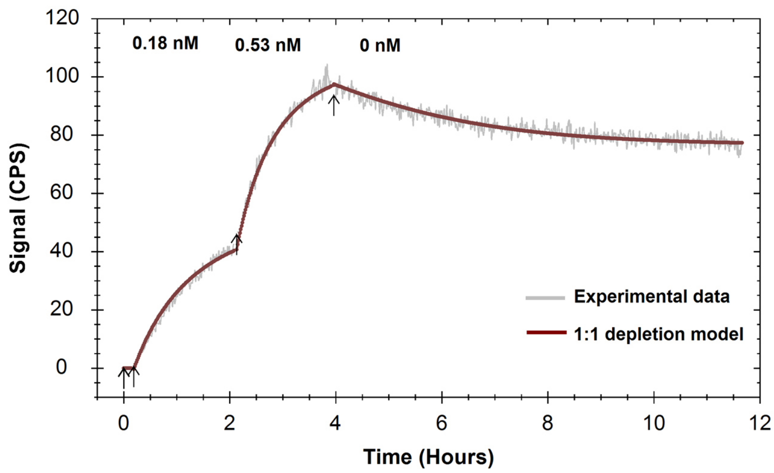


**Figure S2.** Real-time cellular binding of [$^{177}$Lu]Lu-SSO110. Cellular binding curves of [$^{177}$Lu]Lu-SSO110 in high SSTR2-expressing AR42J cells.

1. Castillo Seoane D, De Saint-Hubert M, Ahenkorah S, et al. Gamma counting protocols for the accurate quantification of 225Ac and 213Bi without the need for a secular equilibrium between parent and gamma-emitting daughter. *EJNMMI Radiopharm Chem*. 2022;7:28.
